# The genomic underpinnings of oscillatory biomarkers supporting successful memory encoding in humans

**DOI:** 10.1101/853531

**Authors:** Stefano Berto, Miles Fontenot, Sarah Seger, Fatma Ayhan, Emre Caglayan, Ashwinikumar Kulkarni, Connor Douglas, Carol A. Tamminga, Bradley C. Lega, Genevieve Konopka

## Abstract

In humans, brain oscillations are thought to support critical features of memory formation such as coordination of activity across regions, consolidation, and temporal ordering of events. However, understanding the molecular mechanisms underlining this activity in humans remains a major challenge. Here, we measured memory-sensitive oscillations using direct intracranial electroencephalography recordings from the temporal cortex of patients performing an episodic memory task. By then employing transcriptomics on the resected tissue from the same patients, we linked gene expression with brain oscillations, identifying genes correlated with oscillatory signatures of memory formation across six frequency bands. A co-expression analysis isolated biomarker-specific modules associated with neuropsychiatric disorders as well as ion channel activity. Using single-nuclei transcriptomic data from this resected tissue, we further revealed that biomarker-specific modules are enriched for both excitatory and inhibitory neurons. This unprecedented dataset of patient-specific brain oscillations coupled to genomics unlocks new insights into the genetic mechanisms that support memory encoding. By linking brain expression of these genes to oscillatory patterns, our data help overcome limitations of phenotypic methods to uncover genetic links to memory performance.

## Introduction

Genome-wide association studies and gene expression profiling of the human brain have unlocked the ability to investigate the genetic basis of complex brain phenomena. To date, these datasets have principally been applied to non-invasive imaging studies especially correlations with structural MRI or resting state fMRI^1–5^. Existing methods have relied on published datasets of gene expression from post-mortem brains, meaning that neurophysiological and behavioral data did not come from the same individuals contributing gene expression data^6–8^. This limits the potential impact of such approaches to uncover insights into how genes support key cognitive processes such as episodic memory and highlights the need to develop new datasets in which individuals contribute both neurophysiological and gene expression data^9–11^. Another issue in previous studies is that neurophysiological measurements such as resting state fMRI are not directly linked to cognitive phenomenon. With this in mind, we previously attempted to correlate gene expression levels with oscillatory biomarkers of successful memory encoding^12^, as the fundamental role of these oscillations in supporting memory behavior has been well-established in rodent models and human investigations^13, 14^. These biomarkers are measures of the degree to which memory encoding success modulates oscillatory power in a given frequency band. They were quantified using intracranial electrodes implanted for seizure mapping purposes, with recordings made as participants perform an episodic memory task. In this previous analysis, we used a large database of intracranial EEG (iEEG) recordings obtained over 10 years in order to piece together a distribution of these biomarkers across brain regions. The resulting set of genes that we identified that were correlated with these biomarkers included genes with previously established links to memory formation tested in rodent investigations, genes linked to cognitive disorders such as autism, and novel genes that are prime targets for further investigation. However, as with other studies, this dataset did not have the benefit of both neurophysiological and gene expression information from the same individuals.

With the goal of explicating how gene expression gives rise to brain oscillations and identifying propitious targets for neuromodulation to treat memory disorders, here, we compiled an unprecedented dataset of 16 human subjects who first underwent iEEG during which we measured biomarkers of episodic memory encoding using a well refined signal processing pipeline. These subjects then underwent a temporal lobectomy, during which an en bloc resection of the lateral temporal lobe permitted the acquisition of high quality tissue specimens that were processed immediately upon removal from a common brain region from which in vivo recordings had been previously obtained. These tissue specimens were pathologically normal and we excluded individuals in whom seizure onset included this region.

We made the *a priori* decision to focus on Brodmann area 38 (BA38) in this analysis for the following reasons: 1) the region has been shown to exhibit strong memory related oscillatory biomarkers in multiple investigations^15, 16^; 2) resection of this region is standardized in an en bloc temporal lobectomy operation, allowing the preservation of blood supply to the region until the moment of excision and ensuring a high quality tissue specimen with preservation of cortical architecture for a large sample of tissue; and 3) intracranial EEG investigations preceding temporal lobectomy in this patient population invariably include sampling from this region. Our analysis also offers two additional innovations relative to previous experiments. One is the integration of single nuclei gene expression and chromatin state data using droplet-based microfluidics from surgical human brain samples. Further, we were able to test the cell type-specific expression patterns of genes identified in our analysis as correlated with oscillatory biomarkers using immunofluorescence staining of the related protein products in independent samples, adding confidence to the cell type specificity of the specific genes highlighted.

An inevitable feature of our data set was that all subjects suffered from intractable epilepsy, which presents an important caveat to the interpretations of the results. However, since we examined gene/biomarker correlations across these individuals rather than in comparison to some alternative cohort of data, we were able to institute control methodologies partially accounting for this concern. These included strict artifact rejection routines and the exclusion of data from regions of seizure onset, as well as using matched post-mortem gene expression samples from both unaffected individuals and persons with epilepsy. We explicitly describe these approaches in the Methods and Results.

## Results

### Generation of a within-subjects memory biomarker and gene expression dataset

To determine the relationship between memory-related brain oscillations and gene expression, we analyzed iEEG recorded as subjects encoded episodic memories and gene expression data from the same 16 individuals (see Supplementary Table 1). Biomarkers of successful memory encoding (subsequent memory effects or SMEs), were calculated from recorded iEEG signal by comparing oscillatory patterns during successful versus unsuccessful memory encoding. We used the free recall task, a standard test of episodic memory for which oscillatory patterns have been well-described^17^, and calculated oscillatory biomarkers utilizing our well-established signal processing pipeline^12, 18, 19^ (Fig. 1a and see Methods). Behavioral performance is detailed in Fig. 1. On average, subjects remembered 24.3% of memory items, with a rate of list intrusion of 5.4%. These characteristics are consistent with previous publications of the performance of intracranial EEG subjects on this task^15^.

**Fig 1:**
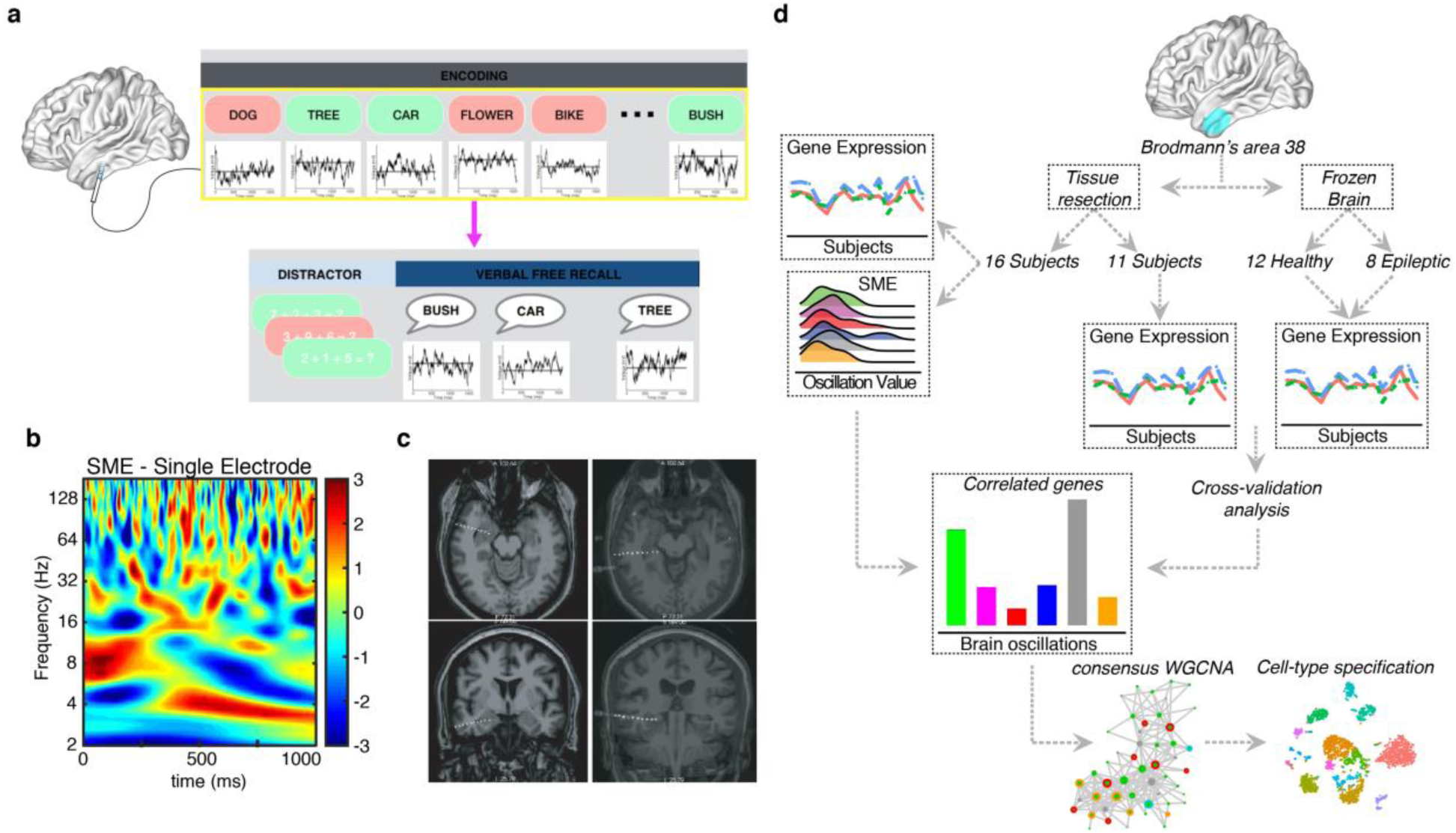
Within-subject study design and quality control. **a,** Schematic of iEEG memory testing. Intracranial electrodes are used to record oscillations as subjects perform an episodic memory task. Oscillatory biomarkers of successful encoding (subsequent memory effects) are calculated by contrasting brain activity recorded as individuals either remembered (green) or forgot (red) each item. **b,** Postoperative MRI after implantation of intracranial electrodes, used for localization. **c,** Example of oscillatory biomarkers of success encoding (subsequent memory effect) recorded from BA38 (full time frequency representation, color axis is z transformed p value for successful/unsuccessful contrast. **d,** Human BA38 RNA-seq from resected tissue was integrated with brain oscillation data derived from subsequent memory effect (SME) analysis, identifying protein-coding genes strongly associated with SMEs (SME genes). Cross-validations with additional human BA38 samples from independent sources was performed. SME genes were prioritized using co-expression networks and specified at the cell-type level using snRNA-seq from BA38.

SMEs were extracted from electrodes located in the temporal pole by first normalizing the iEEG signal following wavelet decomposition and statistically comparing oscillatory values between successful versus unsuccessful encoding events across 56 log spaced frequencies from 2 to 120 Hz. The resulting biomarkers were collapsed into six predefined frequency bands prior to entering these data into our model to estimate gene correlation values (Fig. 1b and see Supplementary Fig. 1a). We observed evidence of SMEs across the frequency spectrum consistent with *a priori* expectation, especially in the gamma frequency range. These 16 subjects then underwent a temporal lobectomy operation. This surgery was performed by a single surgeon (BL) using a technique that was standardized across these patients for obtaining tissue from the anterior temporal pole (BA38) (Fig. 1c). None of the individuals included in the study had gross or radiographic lesions such as temporal sclerosis or cortical dysplasia. Subjects with seizure onset in the temporal pole were excluded.

We generated whole transcriptome RNA-sequencing (RNA-seq) data from the 16 BA38 samples. In addition to the 16 individuals with matched biomarker measurements and gene expression, we generated BA38 RNA-seq data from an additional 11 temporal lobectomies from patients for whom we did not obtain oscillation measurements, and post-mortem tissue from 12 healthy individuals and 8 epileptic patients in order to cross-validate our predictions (Fig. 1d and see Methods). These cross-validated data were used to generate within-subjects correlations of memory biomarker-related brain oscillations and gene expression (Fig. 1d). Principal component analysis revealed that gene expression was uniform across samples with no outliers (see Supplementary Fig. 1b,c). Variance explained by technical and biological covariates was minimal (see Supplementary Fig. 1d) and was removed prior to further analyses. These adjusted gene expression values were then used to calculate gene/biomarker correlations across individuals for each frequency band.

### Memory biomarkers are correlated with gene expression

To determine the relationship between memory biomarkers and gene expression, we used a Spearman’s rank correlation that included the aforementioned cross-validations (see Methods). With this model we identified genes whose expression was correlated with biomarkers in each of the six frequency bands (SME genes). Consistent with *a priori* expectations, we observed significantly correlated genes across the frequency spectrum, with the largest fraction occurring in the gamma ranges (high and low gamma taken together) (Fig. 2a). In line with our previous work^12^, we confirmed beta oscillations as one of the major contributors to gene correlations (n = 769, P < 0.05, *Spearman’s rank correlation*; see Methods and Supplementary Table 2). Surprisingly, we found that delta oscillations showed the greatest number of correlated genes (n = 940, P < 0.05, *Spearman’s rank correlation*) (Fig. 2a and see Supplementary Fig. 2a). To ensure that these observations were specific for a within-subjects study, we used a cross-validation method by repeating the same correlative analysis randomly subsampling from the additional resected and post-mortem brains (see Methods). On average less than 10% of the observed SME genes were identified in the permutations confirming that correlative analysis is largely disrupted when brain oscillations and gene expression are not matched at the same subject level. This result reflects the importance of a within-subject approach to provide a precise association between cognitive recordings and genomics (Fig. 2b; see Supplementary Fig. 2b). Moreover, consistent with our previous results, we noted that there was a substantial sharing of memory biomarker genes across oscillations (Fig. 2c). We found a significant overrepresentation of beta genes in high gamma genes (Fisher’s exact test, FDR corrected, *p* = 1.0×10^-20^, *OR* = 4.58; see Supplementary Fig. 2c). Delta genes were strongly overrepresented in low gamma genes (Fisher’s exact test, FDR corrected, *p* = 5.0×10^-204^, *OR* = 19.3) and in the low-frequency oscillations alpha and theta (Fisher’s exact test, FDR corrected, *p* = 1.0×10^-25^, *OR* = 9.42 and *p* = 4.0×10^-86^, *OR* = 18.7, respectively; see Supplementary Fig. 2c). This pattern specifically suggests that the association of high-frequency power increases and low frequency desynchronization observed across a number of cognitive tasks (but specifically in episodic memory paradigms) may have a common basis in gene expression^20,21^. Data from these 16 individuals also included a control behavioral paradigm in which individuals perform simple mathematical problems, allowing us to observe oscillatory biomarkers linked to this separate cognitive domain (see Methods). We were therefore able to perform the same analysis as above, with the goal of testing whether biomarker— gene correlations were specific for mnemonic processing. Our data set also included cortical thickness estimates for BA38 for each patient extracted from our FreeSurfer^22^ processing routine, allowing us to perform an additional control analysis looking for genes correlated with this measurement. Finally, we looked for gene correlations with memory performance (i.e. behavioral data without regard to any oscillatory biomarker observations). Only a limited number of genes correlated with oscillatory biomarkers overlapped with those identified in these control analyses, reinforcing the unique memory-relevant information obtained by examining gene biomarker correlations (Fig. 2c and see Supplementary Fig. 2d). Genes linked to high gamma oscillations showed the most overlap with math-related genes, which may be consistent with a more general cognitive role for genes linked with high gamma oscillations in this brain region but a more memory-specific role for low gamma and delta correlated genes that did not share enrichment for the math correlated genes.

**Fig. 2:**
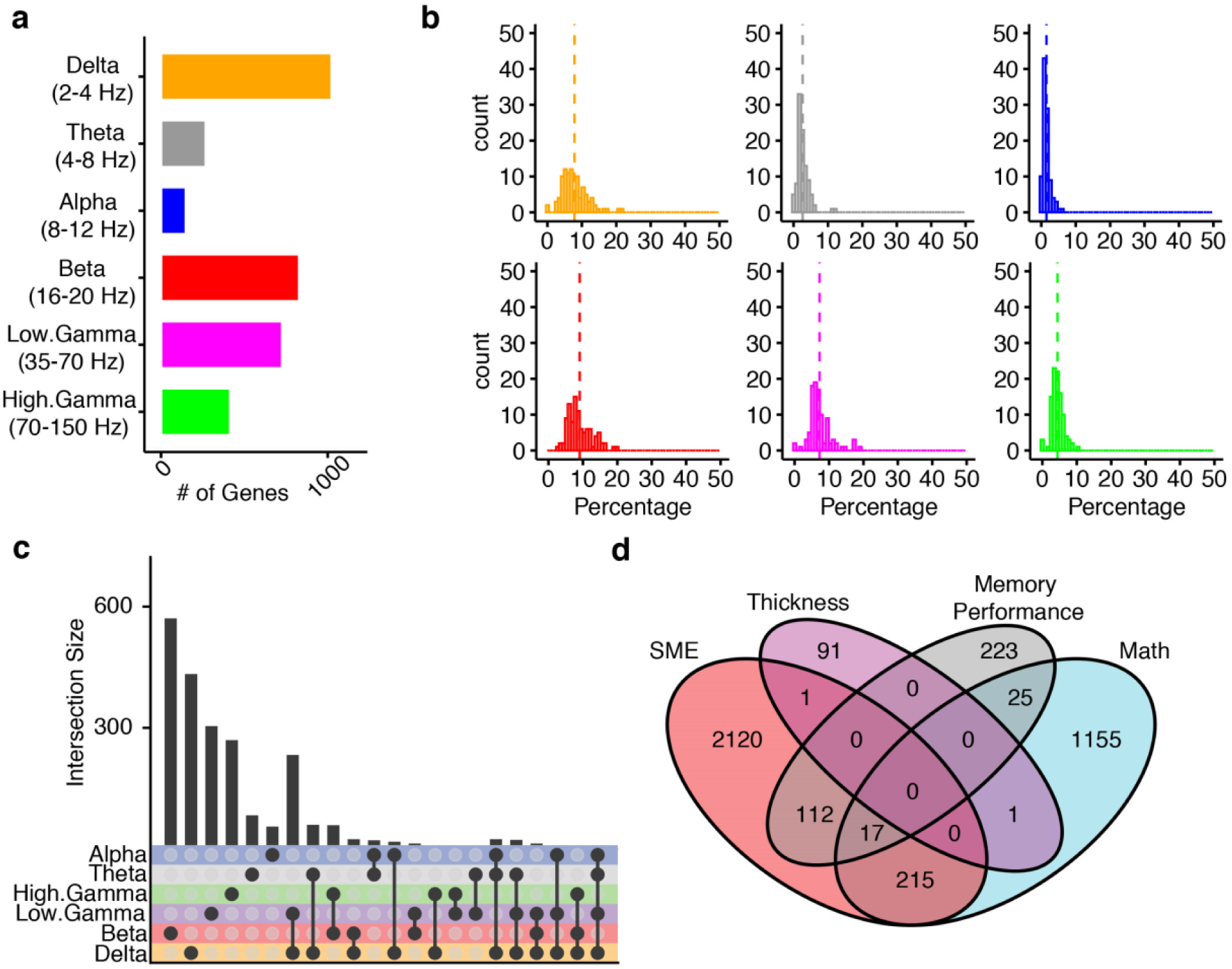
Genes associated with SMEs are distinct. **a,** Bar plot showing the observed number of genes correlated with each brain oscillations. **b**, Histograms showing the percentage of SME genes in the subsampling for each brain oscillation. Dashed line represents the median for the 100 subsampling. **c,** Upset plot indicating shared and specific SME genes between brain oscillations. **d,** Venn diagram showing shared and specific genes between genes associated with memory effect (SME), genes associated with math task (MATH), genes associated with cortical thickness (Thickness), and genes associated with recalled words (Recall).

### Networks refine molecular pathways associated with memory

We sought to understand the functional properties of the genes identified as correlated with oscillatory biomarkers of successful memory encoding. We performed consensus weighted gene co-expression network analysis (consWGCNA^23^; see Methods and Supplementary Table 3) using gene expression from resected temporal lobe tissue together with the post-mortem gene expression datasets. Using this method, we placed the memory genes into a system-level context identifying co-expression networks (e.g. modules of highly correlated genes) linked with brain oscillations to further prioritize genes. We required that identified modules were robust across these multiple expression data sets (see Methods), identifying 26 total modules. Of these, six were significantly associated with biomarker genes (Fig. 3a and see Supplementary Fig. 3b). Two modules were significantly associated with delta oscillations, one module with both delta and low gamma oscillations, and three modules were significantly associated with beta oscillations (Fig. 3a). Notably, we did not detect module association for genes correlated with cortical thickness or recall fraction whereas genes correlated with oscillations during math tasks were associated with an independent module, further confirming that genes associated with memory encoding and their networks are distinct (Fig. 3b). In addition, in three of these modules (WM11, WM12, and WM21), biomarker genes showed higher connectivity compared with the other genes suggesting a central role of these genes in the transcriptome of BA38 (Supplementary Fig. S3c). We also observed convergence of genes and modules associated with biomarker genes from our previous work examining gene/biomarker associations across cortical regions^12^ (Fig. 3c). The convergence of these findings using different patient populations and methods gives confidence to our inferences regarding the link between these genes and mnemonic processes. The two modules positively associated with delta oscillations (WM4 and WM12 respectively) are enriched for genes implicated in ion channel activity (Fig. 3d). Notably, WM4 contains previously identified memory biomarker genes (Fisher’s exact test, FDR corrected; *p* = 0.003, *OR* = 4.4) whereas WM12 is enriched for a previously identified synaptic related biomarker module (Fisher’s exact test, FDR corrected; *p* = 1.0×10^-07^, *OR* = 4.1) (Fig. 3c). *CACNA1C*, one of the hubs of WM12, encodes for a voltage-gated calcium channel associated with risk for schizophrenia, bipolar disorder, major depression disorder and autism^24, 25^. Furthermore, *CACNA1C* has been widely implicated in memory processes and memory retrieval^24–26^. Another WM12 hub is *SHANK2*, a gene encoding a synaptic scaffolding gene. Mutations in *SHANK2* have been linked with autism spectrum disorders, intellectual disability and schizophrenia^27–31^. Moreover, SHANK2 has been associated with learning and memory deficits^32^, further confirming the pivotal role of this WM12 hub gene in memory encoding. Importantly, modules associated with different oscillatory frequency bands exhibited different functional properties. Different than the delta associated modules, modules links to beta oscillations (WM11 and WM22) were significantly associated with alternative splicing and chromatin remodeling (Fig. 3d). In accordance with previous results^12^, we observed that both modules are enriched for genes in SME15, a module linked to beta oscillations with genes implicated in splicing (Fisher’s exact test, FDR corrected; *p* = 2.2×10^-10^, *OR* = 3.9 and *p* = 2.1×10^-05^, *OR* = 3.8 respectively) (Fig. 3c). These data may support alternative splicing regulation as a mechanism for variation in biomarkers observed across individuals. Interestingly, *ATRX*, one of the hub genes of WM11, is important for cognitive development and it is associated with intellectual disability and memory deficits^33, 34^, further supporting the association between chromatin remodeling and alternative splicing with memory^35–37^.

**Fig. 3:**
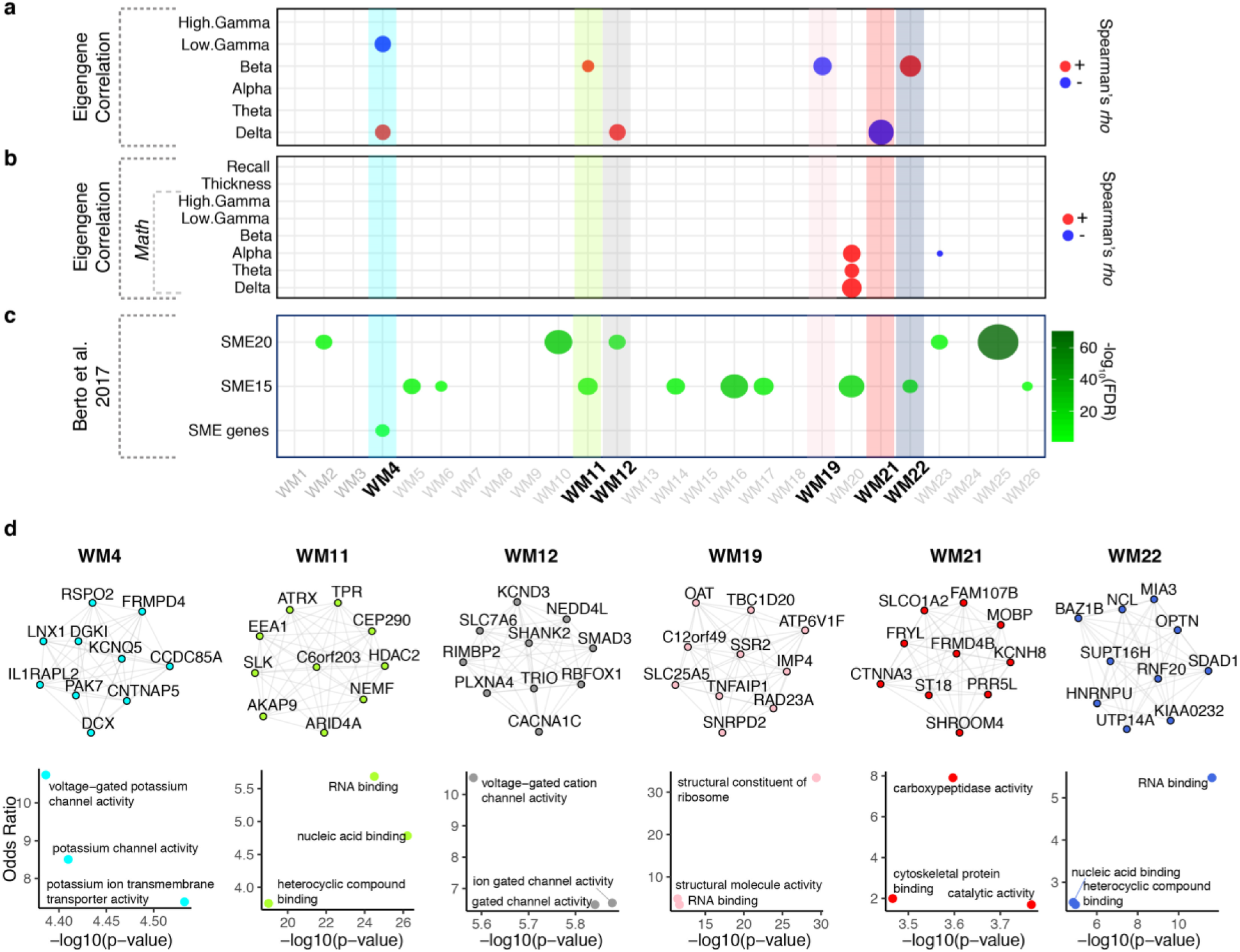
Gene co-expression networks highlight cellular processes implicated in memory encoding. **a,** Bubble-chart showing the module eigengene association with brain oscillations (Spearman’s |*rho*| > 0.5, p < 0.05). **b,** Bubble-chart showing the enrichment for SME genes (Fisher’s exact test, FDR < 0.05). Size represents the - log^10^(FDR). **c,** Bubble-chart showing the enrichment for previously SME genes from a population study^12^. Size represents the -log^10^(FDR). Boldface type indicates the modules associated with brain oscillations. **d,** SME specific modules showing the top 10 hub genes. Edges represent co-expression (*rho*^^2^ > 0.25). Scatterplots represent the top 3 molecular functions of each module. Y-axis = Odds Ratio, X-axis = -log^10^(FDR).

### Modules of memory biomarkers are linked with neuropsychiatric disorders

We next investigated the association of SME modules with genomic data from brain disorders. Using comprehensive transcriptomic and genetic data from multiple disorders (see Methods), we assessed enrichment for genes dysregulated in neuropsychiatric disorders^38^ and GWAS enrichment using LD score regression^39^. The delta associated module WM4 was significantly enriched for down-regulated genes in autism spectrum disorder (ASD; Fisher’s exact test, FDR corrected, *p* = 4.3×10^-4^, *OR* = 2.95) and variants associated with ASD (Fisher’s exact test, FDR corrected, *p* = 0.001) (Fig. 4a,b and see Supplementary Table 4). WM12 showed enrichment for GWAS associated with attention deficit hyperactivity disorder (ADHD; FDR = 0.001), bipolar disorder (BD; FDR = 0.003), major depressive disorder (MDD; FDR = 0.006), schizophrenia (SCZ_2018; FDR = 5.8×10^-6^) and variants associated with educational attainment (FDR = 0.03) and intelligence (FDR = 0.002) (Fig. 4a,b and see Supplementary Table 4). Notably, we found minimal enrichment with variants associated with non-brain related traits and disorders (Supplementary Fig. 4a). We next compared the SME modules with those found in a meta-analysis of transcriptomic data across neuropsychiatric disorders^38^. Both WM4 and WM12 are enriched for a module severely affected in ASD with *RBFOX1* as a predominant hub (geneM1; *p* = 1.5×10^-29^, *OR* = 7.9 and *p* = 2.0×10^-10^, *OR* = 3.92 respectively; Supplementary Fig. 4b). Interestingly, *RBFOX1* is also a hub in WM12, further supporting the role of this gene in neuropsychiatric disorders and memory. The beta module WM11 is enriched for schizophrenia variants (SCZ_2018; FDR = 0.03) (Fig. 4a,b and see Supplementary Table 4) whereas the beta module WM22 is enriched for a splicing module affected in SCZ (geneM19; *p* = 2.6×10^-09^, *OR* = 6.6; see Supplementary Fig. 4b). Overall, the association of delta and beta modules with neuropsychiatric disorders where memory is impaired provide further support to the role of these genes and pathways in episodic memory^40–43^.

**Fig. 4:**
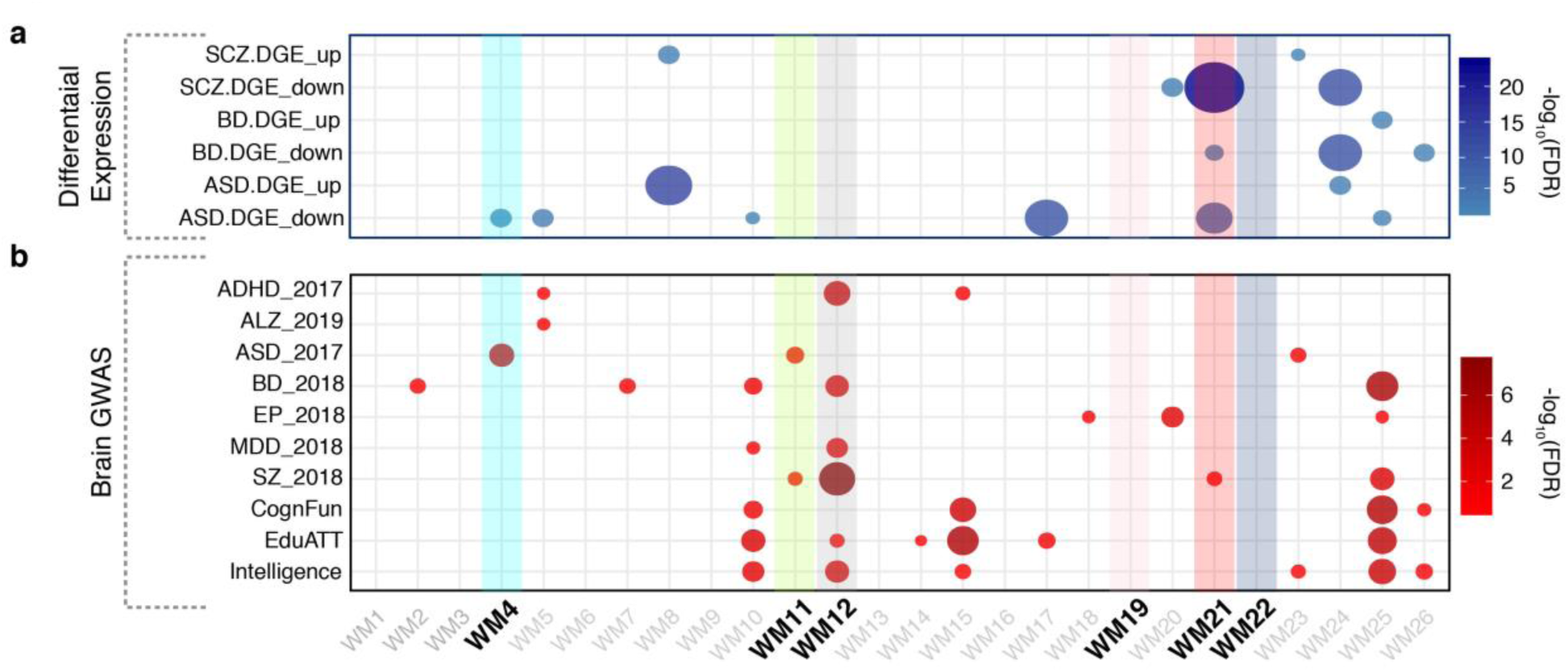
Gene co-expression networks are enriched for neuropsychiatric disorders. **a,** SME-specific modules capture genes dysregulated in neuropsychiatric disorders. Bubble-chart represent the -log^10^(FDR) from a Fisher’s exact enrichment test. Y-axis shows the acronyms for the dysregulated gene data utilized for this analysis. X-axis shows the modules of the present study. **b,** SME-specific modules are enriched for genetic variants associated with neuropsychiatric disorders and cognitive traits. Bubble-chart represent the -log^10^(FDR) from MAGMA. Y-axis shows the acronyms for the GWAS data utilized for this analysis (see Methods). X-axis shows the modules of the present study.

### Modules of memory biomarkers are associated with specific cell-types

To develop cell-type specific associations for the identified correlated genes, we next performed single nuclei RNA-seq (snRNA-seq) analysis on the tissue from three of the original 16 subjects (see Supplementary Table 1). After stringent quality control, expression dimensionality was reduced with principal component analysis (see Methods), and we identified a robust set of 15 transcriptionally-defined clusters (Fig. 5a). To provide confidence to our analysis, we confirmed our clusters with signatures from a previously published snRNA-seq data from the medial temporal gyri^44^ (see Methods and Supplementary Fig. 5a). We defined four inhibitory neuron, eight excitatory neuron and three major non-neuronal clusters. These clusters showed high expression of known major markers for their respective cell types (Fig. 5b and see Supplementary Table 5). We found that the delta modules WM4 and WM12 are strongly enriched for excitatory and inhibitory neurons (Fig. 5c). Interestingly, the module negatively associated with delta, WM21, was enriched for glia cells, with a predominance of oligodendrocyte-related genes (Fig. 5c). Using snRNA-seq from ASD and Alzheimer’s patients^45, 46^, we found that WM4 is significantly enriched for genes dysregulated in Layer 2/3 excitatory neurons in ASD whereas WM21 is significantly enriched for oligodendrocytes markers affected in Alzheimer’s patients (Supplementary Fig. 5b,c). These results further confirm the role of the modules associated with delta oscillations in neuropsychiatric disease at single cell level. WM4 and WM12 are both enriched for delta biomarkers, cognitive-disease related variants, and multiple neuronal types. To further validate our approach for the purpose of future memory neuromodulation strategies specific to brain disorders and cell types, we selected one hub gene from each module for further study. *IL1RAPL2*, an interleukin (IL)-1 receptor accessory protein, is a hub gene in the WM4 module. Intriguingly, along with its paralog IL1RAPL1, IL1RAPL2 promotes functional excitatory synapse and dendritic spine formation^47, 48^ and it is associated with ASD^49^. Our single nuclei RNA-seq data showed IL1RAPL2 has the greatest expression in excitatory neurons but is also expressed in a subset of inhibitory neurons (Fig. 5d). Using fluorescent immunohistochemistry from independently obtained tissue resections, we found that IL1RAPL2 has the greatest overlapping expression with a marker of excitatory neurons, CAMKII, some overlap with a marker of inhibitory neurons, GAD67, and no overlap with a marker of astrocytes, GFAP or a marker of oligodendrocytes, OLIG2 (Fig. 5e,f). Along with the role in excitatory synapse formation, the snRNA-seq and memory biomarkers association indicated that IL1RAPL2 might play an essential role in regulation of memory encoding in humans. Taken together, these results underscore the importance of further studies focused on the role of IL1RAPL2 in memory and excitatory – inhibitory synaptic etiologies.

**Fig. 5:**
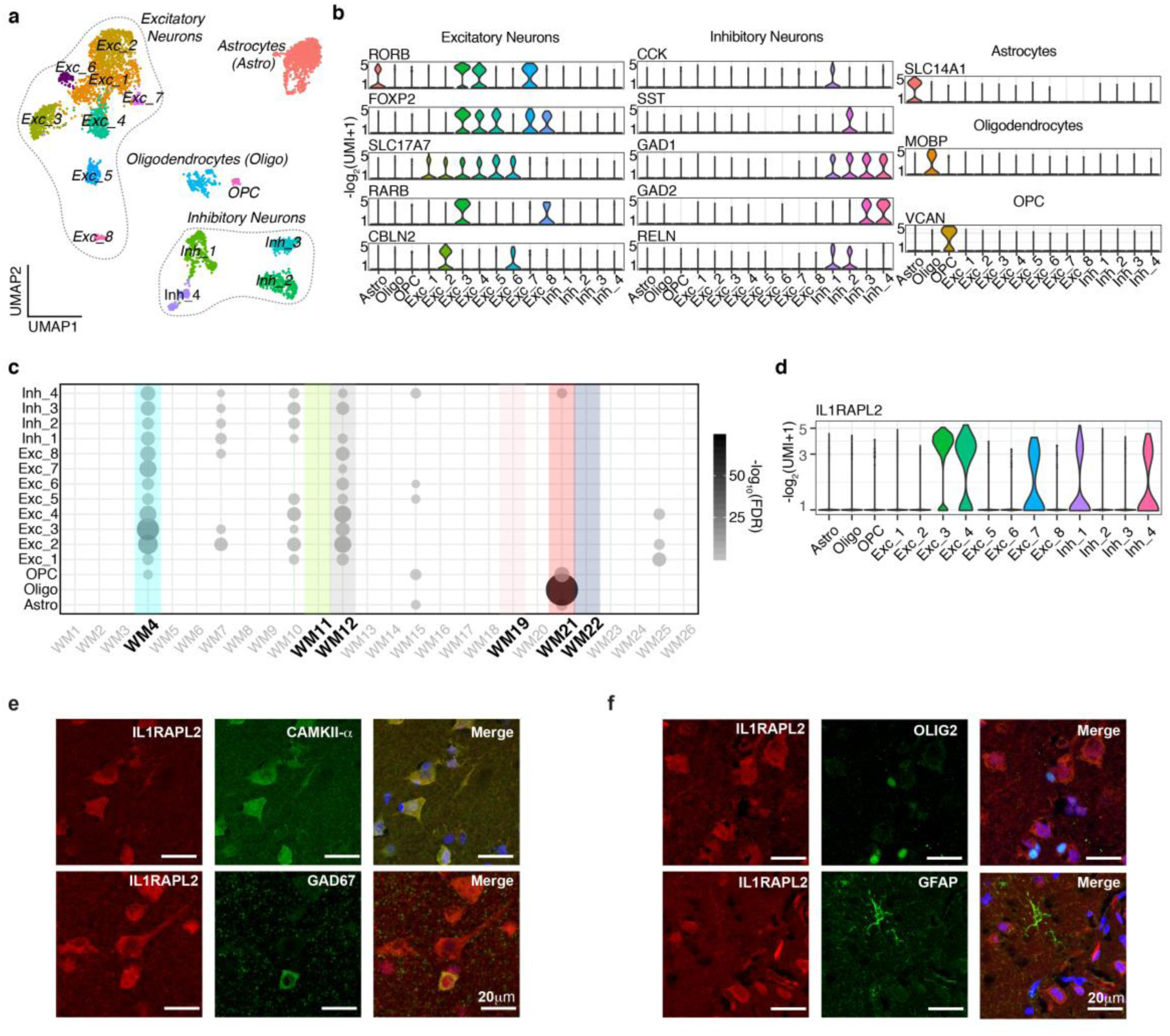
SME-specific modules are enriched for excitatory and inhibitory neurons. **a,** Visualization of the 14 classes of cells using the BA38 snRNA-seq. Cells are displayed based on UMAP. Each dot represents a nucleus. Dashed lines denote the major cell type. *Inh = inhibitory neurons, Exc = excitatory neurons.* **b,** Violin-plots representing gene markers for the main clusters detected. Y-axis represents the -log^2^(UMI). **c,** Cell-type enrichment for SME specific modules. Bubble-chart represents the -log^10^(FDR) from a Fisher’s exact enrichment test. X-axis represents the SME-specific modules. Y-axis represents the cell classes of the present study. Boldface type indicates the modules associated with brain oscillations. **d,** Violin plot representing the expression level (Y-axis) of hub genes of WM4, *IL1RAPL2*. X-axis represents the cell classes of the present study. **e,f,** Immunohistochemistry demonstrate the specific expression of *IL1RAPL2* in excitatory and inhibitory neurons in BA38 (e), not in oligodendrocytes and astrocytes (f).

### Single-nuclei ATAC-seq reveals transcription factors as key regulators of memory-modules

We next sought to understand what transcription factors (TFs) regulate modules of memory biomarkers. To this end, we performed single nuclei ATAC-seq (snATAC-seq) analysis on the tissue from one independent subject (see Supplementary Table 1). After stringent quality controls, we annotated the snATAC-seq clusters using the snRNA-seq (Fig. 6a; see Methods). This multi-omics method allowed us to detect cell-type specific regulatory loci whose accessibility profiles were consistent with the cell-type gene expression. Using motif analysis, we explored the TFs enrichment in the cell-type specific regulatory loci associated with the identified modules of memory biomarkers. Among the modules with cell-type association, motif enrichment was detected only in WM4 and WM12 whereas we did not observe significant enrichment on the oligodendrocytes-specific WM21 (Fig. 6b; see Supplementary table 6). Interestingly, we observed that WM4 genes are highly enriched for Mef2-family motifs, with MEF2A as the highest-scoring motif. MEF2A exhibited enrichment in peaks close to the promoter region of several WM4-hubs suggesting that MEF2A may play a role in the regulation of these memory-linked genes (see Supplementary Fig. 6). Despite the fact that MEF2A showed binding site enrichment, it was not co-expressed with the genes in WM4. We next highlighted the transcription factors that regulate genes in WM12. Interestingly, we found WM12 showed enrichment in motifs of SMAD3, a WM12-hub (Fig. 6b; see Supplementary table 6). Most remarkably, SMAD3 motifs were observed in the promoter region of WM12-hubs associated with neuropsychiatric disorders and memory such as *CACNA1C*^24–26^, *SHANK2*^27–31^, and *NEDD4L*^50–52^ (Fig. 6c). Related to this, previous work has identified NEDD4L as a modulator of SMAD3 turnover during TGFβ signaling transduction^53, 54^. TGFβ signaling plays an important role in modulating neural circuits, axonal formation, synaptic plasticity and cognitive bahaviors^55–58^. Together, the binding site enrichment and co-expression with WM12 hub genes such as NEDD4L indicate the pivotal role of SMAD3 in regulating genes associated with memory oscillations and neuropsychiatric disorders where memory is severely affected. In addition, WM12 contains genes associated with neuronal etiologies, and we found that SMAD3 is specifically expressed in excitatory neurons (Fig. 6d). This result was further confirmed using fluorescent immunohistochemistry from independently obtained tissue resections (Fig. 6e,f). Overall, these results highlight the role of specific transcription factors in the regulation of chromatin landscape necessary to express putative genes associated with memory biomarkers and provide novel molecular entry points for understanding human memory.

**Fig. 6:**
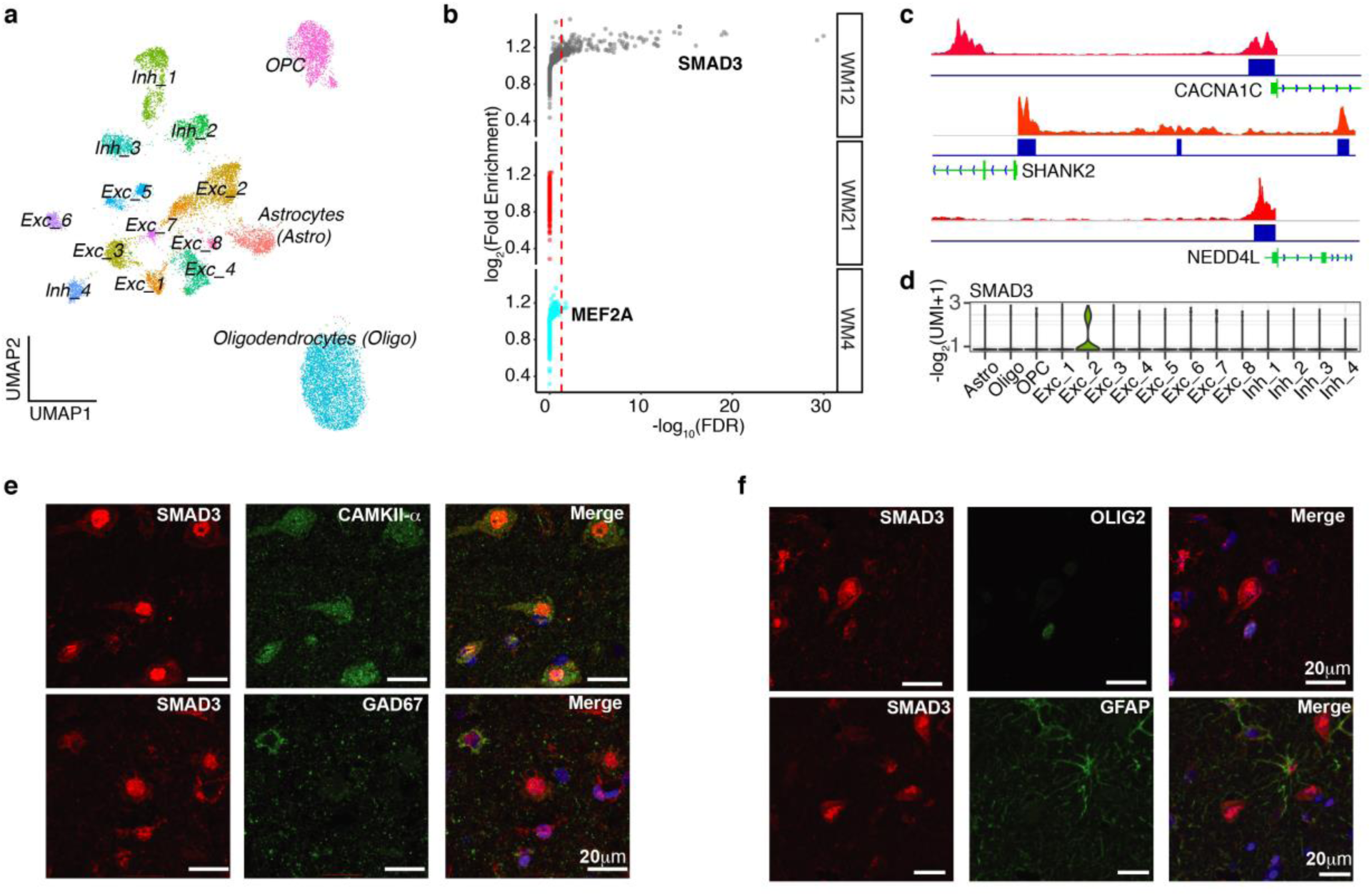
snATAC-seq highlights transcription factors regulating SME-correlated modules. **a,** Visualization of the 14 classes of cells using BA38 snATAC-seq. Cells are displayed based on UMAP. Each dot represents a nucleus. Cell classes were annotated using BA38 snRNA-seq. **b**, Transcription factor binding site enrichment for each of the modules associated with cell-types. MEF2A (WM4) and SMAD3 (WM12) are highlighted. Y-axis represents the -log^2^(Fold Enrichment). X-axis represents the -log^10^(FDR). Dashed line corresponds to FDR = 0.05**. c**, Genomic tracks of snATAC-seq open chromatin regions in the promoters of WM12 hub genes. The SMAD3 binding sites are indicated in blue. **d,** Violin plot representing the expression level (Y-axis) of *SMAD3*, a hub gene of WM12. **e,f,** Immunohistochemistry demonstrates the specific expression of SMAD3 in excitatory (**e**) but not in inhibitory (**e**) neurons, oligodendrocytes (f), and astrocytes (f) in BA38.

## Discussion

We set out to understand the genomic underpinnings of brain oscillations that support episodic memory encoding in humans, with the ultimate goal of identifying genes that are propitious targets for neuromodulation strategies to treat memory disorders. Using an unparalleled data set of 16 human subjects from which we obtained measurements of brain oscillations linked to successful episodic memory encoding as well as whole transcriptomic data from the temporal pole *in the same individuals*, we identified modules of genes that link specific cell types and cellular functions with memory biomarkers. One point worthy of emphasis is that our analysis is fundamentally different than previous attempts to correlate gene expression with behavioral measurements such as memory performance^59–61^. Oscillatory correlates of successful memory encoding represent an “intermediate step” between gene regulation and memory behavior. Oscillations are localized to the brain region in which they are recorded using intracranial depth electrodes and are dissociable into frequency bands with distinct properties. Linking neurophysiological measurements (such as these oscillatory biomarkers) with gene expression data will establish specific testable hypotheses for subsequent investigations for these genes we have identified. Our work sheds light on the molecular mechanisms that give rise to oscillatory correlates of successful memory encoding^62^. In this regard, our observation that delta oscillations are linked to ion channel genes and further that these genes tend to be expressed in oligodendrocytes leads to the fascinating implication that the generation of low-frequency oscillatory patterns linked to mnemonic processing in humans is at least partially dependent upon glial modulation of oscillations. This is based upon a combination of our observations across all subjects as well as the single nuclei expression analysis. This conclusion is supported by the role of oligodendrocytes in learning and memory acting on depolarization of membrane potential^63–66^, which accelerates axonal conduction and ion channel activity as reflected by the delta modules with positive association (WM4 and WM12). We note that we observed interesting properties for genes correlated with delta oscillations, but not theta oscillations which prima fascia runs contrary to rodent data that universally implicate theta frequency activity in successful memory formation^67^. However, in the human temporal lobe, oscillations outside of the 4-9 Hz range routinely exhibit memory—relevant properties, including cross-frequency coupling, and as such our findings are in line with previous observations using biomarkers of successful memory encoding in humans^68, 69^. In humans these low frequency oscillations represent a consistent feature of oscillatory signatures of memory formation, including influence on the timing of single unit activity^68, 70–72^. The significant representation of delta correlated genes in our analysis may reflect the functional importance of these low frequency components in humans. One necessary caveat in interpreting our data is the fact that all subjects suffer from intractable epilepsy. Clearly, the use of such a subject population is necessary to generate these highly valuable data with both in vivo oscillations and gene expression data from the same individuals in humans. However, several features of our analysis such as under-enrichment for genetic variants associated with epilepsy and the data integration with epileptic and healthy tissues give us confidence that the insights we have uncovered represent more generalizable associations between gene expression and brain oscillations. Further, numerous human studies have established that observations in iEEG patients have correlates using noninvasive studies in normal control individuals and in animal models^73–76^. Additionally, we employed strict artifact rejection criteria and eliminated electrodes located in the seizure onset zone in our analysis, reducing the impact of abnormal activity on observed oscillatory biomarkers^76, 77^. We also included several control steps in our analysis including incorporation of duration of epilepsy to adjust gene expression values. Finally, several of the key genes we have identified (for example *CACNA1C*, *MET*, and *SMAD3*) have been independently shown to be linked to memory processing in data from non-epileptic individuals. Collectively, this translational work establishes an experimental and analytical approach for deconstructing human behavioral and cognitive traits such as memory using integrative physiological and multi-omics techniques. Integration of single nucleus transcriptomic and epigenomic data allowed us to identify the cell type specificity of the memory-related gene co-expression modules as well as potential regulators of these modules (e.g. SMAD3and MEF2A). For example, our data suggest that SMAD3 acts as a regulator of genes and molecular pathways involved in memory encoding-related delta oscillations, specifically in excitatory neurons. This molecular characterization of human memory highlights a key regulator that can be further studied in model systems. We anticipate that this within-subjects approach can be used in future studies to highlight molecular pathways of other human complex traits with the ultimate goal of identifying therapeutic targets and linking clinical and genomic data at the individual subject level. Importantly, investigations using animal and in vitro models will be necessary to definitively characterize the memory related properties of the genes identified in our analysis.

## Methods

### EXPERIMENTAL MODEL AND SUBJECT DETAILS

#### Resected brain samples

All surgical samples included in this study were BA38 resections from patients with temporal lobe epilepsy. The brain specimen was dropped into ice-cold 1X PBS in a 50mL conical tube immediately after removal from the patient. After 4-5 inversions, the tissue sample was transferred to a fresh tube with ice-cold 1X PBS for a second wash. The specimen was then moved to a petri dish and dissected grossly by scalpel into ∼12 sub-samples and frozen immediately in individual Eppendorf tubes in liquid nitrogen as the tubes were filled. Care was taken to avoid major blood vessels. Gray matter was prioritized over tracts of white matter in an attempt to increase homogeneity and consistency of results across all samples. Time from removal of brain to flash freezing ranged from roughly two minutes for the first piece to about seven minutes for the last sub-sample. Three to four of the sub-samples were extracted for RNA, and the sub-sample with the highest RIN value was selected for RNA-sequencing. See Supplementary Table S1 for detailed demographic information.

#### Postmortem brain samples

Twelve samples of BA38 were obtained from the Dallas Brain Collection. These tissue samples were donated from individuals without a history of neurological or psychiatric disorders as previously published^78^. Eight samples of BA38 were obtained from the University of Maryland Brain and Tissue Bank. These samples were donated from individuals with epilepsy. See Supplementary Table S1 for detailed demographic information.

#### RNA-sequencing (RNA-seq)

Total RNA was purified using an miRNeasy kit (#217004, Qiagen) following the manufacturer’s recommendations. RNA-seq libraries from mRNA were prepared in-house as previously described^79^. Sequencing was performed on randomly pooled samples by the McDermott Sequencing Core at UT Southwestern on an Illumina NextSeq 500 sequencer. Single-end, 75-base-pair (bp) reads were generated.

#### Isolation of nuclei from resected brain tissues (snRNA-seq)

Nuclei were isolated as previously described^80^ https://www.protocols.io/view/rapid-nuclei-isolation-from-human-brain-scpeavn. Surgically resected cortical tissue was homogenized using a glass dounce homogenizer in 2 ml of ice-cold Nuclei EZ lysis buffer (#EZ PREP NUC-101, Sigma) and was incubated on ice for 5 min. Nuclei were centrifuged at 500 × g for 5 min at 4 °C, washed with 4 ml ice-cold Nuclei EZ lysis buffer and, incubated on ice for 5 min. Nuclei were centrifuged at 500 × g for 5 min at 4 °C. After centrifugation, the nuclei were resuspended in 1 ml of nuclei suspension buffer (NSB) consisting of 1XPBS, 1%BSA (#AM2618, Thermo Fisher Scientific) and 0.2U/ul RNAse inhibitor (#AM2694, Thermo Fisher Scientific) and were filtered through a 40-μm Flowmi Cell Strainer (#H13680-0040, Bel-Art). Nuclei concentration was determined using 0.4% Trypan Blue (#15250061, Thermo Fisher Scientific). Final concentration of 1000 nuclei/μl was adjusted with NSB. Droplet-based snRNA-seq libraries were prepared using the Chromium Single Cell 3’ v2 (#120237, 10x Genomics) according to the manufacturer’s protocol^81^. Libraries were sequenced using an Illumina NextSeq 500 at the McDermott Sequencing Core at UT Southwestern.

#### Isolation of nuclei from resected brain tissue (snATAC-seq)

For snATAC-seq, nuclei were isolated as described above. After lysis, the nuclei were washed once in 500 μl of nuclei wash buffer consisting of 10mM Tris-HCl (pH 7.4), 10mM NaCl, 3mM MgCl2, 1% BSA and, 0.1% Tween-20. Nuclei were resuspended in 500 μl of 1X Nuclei Buffer (10X Genomics). Debris was removed with a density gradient centrifugation using the Nuclei PURE 2M Sucrose Cushion Solution and Nuclei PURE Sucrose Cushion Buffer from Nuclei PURE Prep Isolation Kit (#NUC201-1KT, Sigma Aldrich). Nuclei PURE 2M Sucrose Cushion Solution and Nuclei PURE Sucrose Cushion Buffer were first mixed in 9:1 ratio. 500 μl of the resulting sucrose buffer was added to a 2 ml Eppendorf tube. 900 μl of the sucrose buffer was added to 500 μl of isolated nuclei in NSB. 1400 μl nuclei suspension was layered to the top of sucrose buffer. This gradient was centrifuged at 13, 000 x g for 45 min at 4 °C. Nuclei pellets were resuspended and washed once in nuclei wash buffer. Nuclei concentration and integrity were determined using Ethidium Homodimer-1 (EthD-1) (#E1169, Thermo Fisher Scientific). Finally, nuclei were resuspended in 1X Nuclei Buffer at concentration of 4000 nuclei/μl for single-cell ATAC-sequencing. Droplet-based single-cell ATACseq libraries were prepared using the Chromium Single Cell ATAC Kit Solution v1.0 (10X Genomics) and following the Chromium Single Cell ATAC Reagent Kits User Guide: CG000168 Rev B. The library was sequenced using an Illumina NextSeq 500 at the McDermott Sequencing Core at UT Southwestern.

#### Immunofluorescence staining of human tissue

Fresh surgically resected tissue was fixed in 4% PFA in 1x PBS 24-48h at 4C and then cryoprotected in a 30% sucrose solution. The tissue was sectioned at 7 µm using a cryostat (Leica). Sections underwent heat induced antigen retrieval in a citrate buffer (pH 6.0) for 10 min at 95 C. Sections were blocked with 2% fetal bovine serum (FBS) in 0.1M Tris (pH 7.6) for 1hour at room temperature. After blocking, the sections were incubated with primary antibodies in 0.1 M Tris pH 7.6/2% FBS overnight at 4C and subsequently incubated with secondary antibodies in 0.1M Tris pH pH 7.6/2% FBS for 1 h at room temperature. Sections were immersed in 0.25% Sudan Black solution to quench lipofuscin auto-fluoresce and counterstained with 4′-6-diamidino-2-phenylindole (DAPI). Sections were mounted and cover slipped using ProLong Diamond Antifade Mountant (#P36970, Thermo Fisher Scientific). The following antibodies and dilutions were used: goat α-IL1RAPL2 (#PA5-47039, Thermo Fisher Scientific 1:20), rat α-SMAD3 (#MAB4038, R&D Systems, 1:100), rabbit α-CaMKII alpha (#PA514315, Thermo Fisher Scientific, 1:50), chicken α-GFAP (#ab4674, Abcam, 1:400), mouse α-GAD67 (#MAB5406, Millipore, 1:200), mouse α-OLIG2 (#MABN50,Millipore, 1:200), species-specific secondary antibodies produced in donkey and conjugated to Alexa Fluor 488, Alexa Fluor 555, or Alexa Fluor 647 (Thermo Fisher Scientific, 1:800). Images were acquired using a 63X oil objective on a Zeiss LSM 880 confocal microscope.

## COMPUTATIONAL METHODS

### RNA-seq mapping, QC and expression quantification

Reads were aligned to the human hg38 reference genome using STAR 2.5.2b^82^ with the following parameters: “*--outFilterMultimapNmax 10 --alignSJoverhangMin 10 -- alignSJDBoverhangMin 1 --outFilterMismatchNmax 3 --twopassMode Basic*”. For each sample, a BAM file including mapped and unmapped reads that spanned splice junctions was produced. Secondary alignment and multi-mapped reads were further removed using in-house scripts. Only uniquely mapped reads were retained for further analyses. Quality control metrics were performed using RSeQC^83^ using the hg38 gene model provided. These steps include: number of reads after multiple-step filtering, ribosomal RNA reads depletion, and defining reads mapped to exons, UTRs, and intronic regions. Picard tool was implemented to refine the QC metrics (http://broadinstitute.github.io/picard/). Gencode annotation for hg38 (version 24) was used as reference alignment annotation and downstream quantification. Gene level expression was calculated using HTseq version 0.9.1^84^ using intersection-strict mode by gene. Counts were calculated based on protein-coding genes from the annotation file.

### Covariate adjustment

Counts were normalized using counts per million reads (CPM) with *edgeR* package in R ^85^. Normalized data were log2 scaled with an offset of 1. Genes with no reads were removed. Normalized data were assessed for effects from known biological covariates (*Sex*, *Age, Race, Ethnicity, Hemisphere, Epilepsy Duration*), technical variables related to sample processing (RNA integrity number: *RIN, Batch*). Other biological and technical covariates (*Post-mortem interval: PMI*) were not considered for the analysis because they were confounded with the brain resected data. The data were adjusted for technical covariates using a linear model:

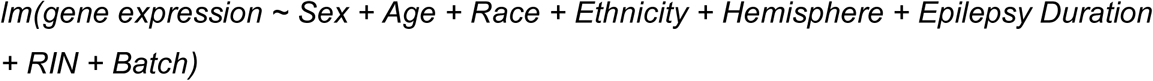

Adjusted CPM values were used for SME correlation, co-expression analysis and visualization.

### Correlation analysis and cross-validation analysis

Spearman’s rank correlation was performed between each of the 6 memory brain oscillations and gene expression. We also utilized this method for 6 math brain oscillations, thickness and behavioral performances.

For this analysis we used:

1. *Within Subject: bulk RNA-seq from Brodmann’s area 38 resected tissue of 16 subjects with calculated SME (WrS).*

We next performed cross-validation analysis using data from:

1. *Additional Subjects: additional 11 subjects of bulk RNA-seq from Brodmann’s area 38 resected tissue without SME (ArS).*
2. *Independent Data Healthy: bulk RNA-seq from Brodmann’s area 38 frozen tissue of 12 subjects (HfS).*
3. *Independent Data Epilepsy: bulk RNA-seq from Brodmann’s area 38 frozen tissue of 8 subjects with Epilepsy (EfS).*

Bootstrap was applied randomly sampling 16 subjects (as WrS) from the composite data and recalculating the correlation 100 times. We then calculated a Monte Carlo p-value comparing the observed effect with the simulated effects for each gene by:

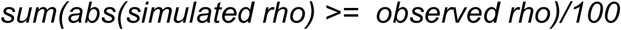

We calculated two *Monte Carlo* p-values:

1. *BootP based on WrS + ArS (only resected tissues)*
2. *BootP_All based on WrS + ArS + HfS + EfS (resected tissues and frozen tissues)* We additionally applied a permutation approach shuffling the gene expression of WS and recalculating 100 times the correlation between oscillations and gene expression. We then calculated a *Monte Carlo* p-value (PermP). Nominal p.value < 0.05, PermP < 0.05, BootP < 0.05 and BootP_All < 0.05 were used to filter for significant correlations as reported in Table S2. Finally, we used *ArS + HfS + EfS* (additional resected tissues and frozen tissues) for the subsampling showed in the Figure 2b and Supplementary Fig. 2b. Briefly, we subsample from *ArS + HfS + EfS* 16 random subjects 100 times and performed the correlative analysis. We next selected the number of genes that were significant in each of the 100 shuffle and compared them with the observed SME genes.

### Co-expression network analysis

To identify modules of co-expressed genes in the RNA-seq data, we carried out weighted gene co-expression network analysis (WGCNA)^23^. We applied a consensus analysis based on WrS + ArS + HfS + EfS data defining modules highly preserved across multiple datasets. This method was applied to reduce the potential noise between different types of data. A soft-threshold power was automatically calculated to achieve approximate scale-free topology (R^2^>0.85). Networks were constructed with *blockwiseConsensusModules* function with biweight midcorrelation (bicor). We used *corType = bicor, networkType = signed, TOMtype = signed, TOMDenom = mean, maxBlockSize = 16000, mergingThresh = 0.15, minCoreKME = 0.5, minKMEtoStay = 0.6, reassignThreshold = 1e-10, deepSplit = 4, detectCutHeight = 0.999, minModuleSize = 50*. The modules were then determined using the dynamic tree-cutting algorithm. To ensure robustness of the observed network, we used a permutation approach recalculating the networks 200 times and comparing the observed connectivity per gene with the randomized one. None of the randomized networks showed similar connectivity, providing robustness to the network inference. Module sizes were chosen to detect small modules driven by potential noise on the adjusted data. Deep split of 4 was used to more aggressively split the data and create more specific modules. Spearman’s rank correlation was used to compute module eigengene – memory biomarkers associations.

### Single-nuclei RNA-seq analysis

Single-nuclei RNA-seq data from BA38 was processed using *mkfastq* command from 10X Genomics CellRanger v3.0.1. Extracted paired-end fastq files (26 bp long R1 - cell barcode and UMI sequence information, 124 bp long R2 - transcript sequence information) were checked for read quality using FastQC. Gene counts were obtained by aligning reads to the hg38 genome using an in-house pipeline. UMI-tools^86^ were used to match barcode and reads. Reads were aligned to the human hg38 reference genome using STAR 2.5.2b^82^. Gencode annotation for hg38 (version 24) was used as reference alignment annotation. Gene level expression was calculated using featureCounts^87^ by gene. UMIs were further calculated using UMI-tools. Seurat^88^ standard pipeline was used for downstream analysis and cell-markers definition (see Code Availability).

### Single-nuclei ATAC-seq analysis

Single-nuclei ATAC-seq data from BA38 was processed using Cell Ranger ATAC pipeline (see Code Availability). Seurat extension Signac^88^ was used for additional filtering, clustering and annotation. Cells with total fragments in peaks less than 1000 or less than 15% of the total fragments were not considered for future analysis. Clustering and creating a gene activity matrix were done with the default parameters. Only the cells with >0.5 confidence in annotation were considered for downstream analysis. Motif enrichment was tested for the upstream regions of the genes in each module. Motif analysis was performed only for the modules with cell-type enrichment (WM4/WM12: excitatory-inhibitory clusters, WM21: oligodendrocyte-OPC clusters). Fragments for excitatory-inhibitory clusters and oligodendrocyte-OPC clusters were extracted separately from Cell Ranger’s fragments.tsv file. For each cut site, the fragments.tsv file was adjusted to contain 200bp around the cut site and peaks were called using MACS2^89^. The CIS-BP database for human was used for enrichment^90^. Only TFs with directly determined motifs were kept. TFs were filtered for presence in > 30% of cells in the cluster that is tested for enrichment. A motif matrix (peaks in rows, motifs in columns) was created with CreateMotifMatrix from Signac. Using the FindMotifs function from Signac, enrichment of each TF was tested for the upstream peaks of module genes versus upstream peaks of all genes using all the peaks as background. Peak visualization was done using IGV^91^.

### Functional Enrichment

The functional annotation of differentially expressed and co-expressed genes was performed using ToppGene^92^. We used GO and KEGG databases. Pathways containing between 5 and 2000 genes were retained. A Benjamini-Hochberg FDR (*P* < 0.05) was applied as a multiple comparisons adjustment.

### GWAS data and enrichment

Summary statistics for GWAS studies on neuropsychiatric disorders and non-brain disorders were downloaded from Psychiatric Genomics Consortium and GIANT Consortium^93–107^. MAGMA v1.04^39^ was used for gene set analysis. Supplementary Table 4 reports MAGMA statistics for each of the GWAS data analyzed. GWAS acronyms were used for the figures (e.g. *ADHD = attention deficit hyperactivity disorder, ASD = autism spectrum disorders, ALZ = Alzheimer’s disease, BIP = bipolar disorder, EP = epilepsy, MDD = major depressive disorder, SZ = schizophrenia*, *EduAtt = educational attainment, Intelligence = Intelligence, CognFunc = cognitive functions, BMI = body mass index, CHD = coronary artery disease, DIAB = diabetes, HGT = height, OSTEO = osteoporosis)*.

### Gene set enrichment

Gene set enrichment was applied to correlated genes and SME genes^12^ from our previous study as shown in Fig. 3c, neuropsychiatric DEGs as shown in Fig. 4a and Supplementary Fig. 4b, and cell-type markers as shown in Fig. 5c and Supplementary Fig. 5. We used a Fisher’s exact test in R with the following parameters: alternative = “greater”, conf.level = 0.95. We reported Odds Ratios (OR) and Benjamini-Hochberg adjusted P-values (FDR).

## Acknowledgments

We thank the patients for participating in the study and the donors and their families for the additional tissue samples. We also thank Kelly Gleason for assistance with post-mortem samples. G. K. is a Jon Heighten Scholar in Autism Research at UT Southwestern. This work was supported by NIMH (F30MH105158) to M.F.; NIDA (5T32DA007290-25) and NHBLI (1T32HL139438-01A1) to F.A.; NINDS (NS106447), a UT BRAIN Initiative Seed Grant (366582), the Chilton Foundation, and the National Center for Advancing Translational Sciences of the NIH under the Center for Translational Medicine’s award number UL1TR001105, to B.L. and G.K.; NINDS (NS107357) to B.L.; and NIMH (103517), The Chan Zuckerberg Initiative, an advised fund of Silicon Valley Community Foundation (HCA-A-1704-01747), and the James S. McDonnell Foundation 21^st^ Century Science Initiative in Understanding Human Cognition – Scholar Award (220020467) to G.K.. Post-mortem human tissue samples were obtained from the NIH NeuroBioBank (The Harvard Brain Tissue Resource Center, funded through HHSN-271-2013-537 00030C; the Human Brain and Spinal Fluid Resource Center, VA West Los Angeles Healthcare Center; and the University of Miami Brain Endowment Bank) and the UT Neuropsychiatry Research Program (Dallas Brain Collection).

## Author Contributions

S.B., B.L., and G.K. analyzed the data and wrote the paper. M.F. and C.D. collected surgical samples, processed RNA and generated bulk RNA-seq libraries. F.A. generated snRNA-seq/ATAC-seq data and performed IHC. A.K. pre-processed the snRNA-seq data. E.C. pre-processed the snATAC-seq data. C.T. provided post-mortem human brain tissue. S.S. analyzed the oscillation data. B.L. conducted all surgical procedures and memory testing. B.L. and G.K. designed and supervised the study, and provided intellectual guidance. All authors discussed the results and commented on the manuscript.

## References

1 Morgan, S. E. et al. Cortical patterning of abnormal morphometric similarity in psychosis is associated with brain expression of schizophrenia-related genes. Proc Natl Acad Sci U S A 116, 9604–9609, doi:10.1073/pnas.1820754116 (2019).

2 Lombardo, M. V. et al. Large-scale associations between the leukocyte transcriptome and BOLD responses to speech differ in autism early language outcome subtypes. Nat Neurosci 21, 1680–1688, doi:10.1038/s41593-018-0281-3 (2018).

3 Romero-Garcia, R., Warrier, V., Bullmore, E. T., Baron-Cohen, S. & Bethlehem, R. A. I. Synaptic and transcriptionally downregulated genes are associated with cortical thickness differences in autism. Mol Psychiatry 24, 1053–1064, doi:10.1038/s41380-018-0023-7 (2019).

4 Pantazatos, S. P. & Li, X. Commentary: BRAIN NETWORKS. Correlated Gene Expression Supports Synchronous Activity in Brain Networks. Science 348, 1241–4. Front Neurosci 11, 412, doi:10.3389/fnins.2017.00412 (2017).

5 Wang, G. Z. et al. Correspondence between Resting-State Activity and Brain Gene Expression. Neuron 88, 659–666, doi:10.1016/j.neuron.2015.10.022 (2015).

6 Patania, A. et al. Topological gene expression networks recapitulate brain anatomy and function. Netw Neurosci 3, 744–762, doi:10.1162/netn_a_00094 (2019).

7 Le, B. D. & Stein, J. L. Mapping causal pathways from genetics to neuropsychiatric disorders using genome-wide imaging genetics: Current status and future directions. Psychiatry Clin Neurosci 73, 357–369, doi:10.1111/pcn.12839 (2019).

8 Smit, D. J. A. et al. Genome-wide association analysis links multiple psychiatric liability genes to oscillatory brain activity. Hum Brain Mapp 39, 4183–4195, doi:10.1002/hbm.24238 (2018).

9 Konopka, G. Cognitive genomics: Linking genes to behavior in the human brain. Netw Neurosci 1, 3–13, doi:10.1162/NETN_a_00003 (2017).

10 Udden, J. et al. Towards robust functional neuroimaging genetics of cognition. J Neurosci, doi:10.1523/JNEUROSCI.0888-19.2019 (2019).

11 Mufford, M. S. et al. Neuroimaging genomics in psychiatry-a translational approach. Genome Med 9, 102, doi:10.1186/s13073-017-0496-z (2017).

12 Berto, S., Wang, G. Z., Germi, J., Lega, B. C. & Konopka, G. Human Genomic Signatures of Brain Oscillations During Memory Encoding. Cereb Cortex 28, 1733–1748, doi:10.1093/cercor/bhx083 (2018).

13 Long, N. M., Burke, J. F. & Kahana, M. J. Subsequent memory effect in intracranial and scalp EEG. Neuroimage 84, 488–494, doi:10.1016/j.neuroimage.2013.08.052 (2014).

14 Mukamel, R. & Fried, I. Human intracranial recordings and cognitive neuroscience. Annu Rev Psychol 63, 511–537, doi:10.1146/annurev-psych-120709-145401 (2012).

15 Sederberg, P. B. et al. Hippocampal and neocortical gamma oscillations predict memory formation in humans. Cereb Cortex 17, 1190–1196, doi:10.1093/cercor/bhl030 (2007).

16 Nakamura, K. & Kubota, K. The primate temporal pole: its putative role in object recognition and memory. Behav Brain Res 77, 53–77, doi:10.1016/0166-4328(95)00227-8 (1996).

17 Sederberg, P. B., Howard, M. W. & Kahana, M. J. A context-based theory of recency and contiguity in free recall. Psychol Rev 115, 893–912, doi:10.1037/a0013396 (2008).

18 Arora, A. et al. Comparison of logistic regression, support vector machines, and deep learning classifiers for predicting memory encoding success using human intracranial EEG recordings. J Neural Eng 15, 066028, doi:10.1088/1741-2552/aae131 (2018).

19 Lin, J. J. et al. Theta band power increases in the posterior hippocampus predict successful episodic memory encoding in humans. Hippocampus 27, 1040–1053, doi:10.1002/hipo.22751 (2017).

20 Ezzyat, Y. et al. Direct Brain Stimulation Modulates Encoding States and Memory Performance in Humans. Curr Biol 27, 1251–1258, doi:10.1016/j.cub.2017.03.028 (2017).

21 Hanslmayr, S., Staresina, B. P. & Bowman, H. Oscillations and Episodic Memory: Addressing the Synchronization/Desynchronization Conundrum. Trends Neurosci 39, 16–25, doi:10.1016/j.tins.2015.11.004 (2016).

22 Fischl, B. FreeSurfer. Neuroimage 62, 774–781, doi:10.1016/j.neuroimage.2012.01.021 (2012).

23 Langfelder, P. & Horvath, S. WGCNA: an R package for weighted correlation network analysis. BMC Bioinformatics 9, 559, doi:10.1186/1471-2105-9-559 (2008).

24 Sykes, L., Clifton, N. E., Hall, J. & Thomas, K. L. Regulation of the Expression of the Psychiatric Risk Gene Cacna1c during Associative Learning. Mol Neuropsychiatry 4, 149–157, doi:10.1159/000493917 (2018).

25 Yoshimizu, T. et al. Functional implications of a psychiatric risk variant within CACNA1C in induced human neurons. Mol Psychiatry 20, 162–169, doi:10.1038/mp.2014.143 (2015).

26 Cosgrove, D. et al. Cognitive Characterization of Schizophrenia Risk Variants Involved in Synaptic Transmission: Evidence of CACNA1C’s Role in Working Memory. Neuropsychopharmacology 42, 2612–2622, doi:10.1038/npp.2017.123 (2017).

27 Berkel, S. et al. Mutations in the SHANK2 synaptic scaffolding gene in autism spectrum disorder and mental retardation. Nat Genet 42, 489–491, doi:10.1038/ng.589 (2010).

28 Berkel, S. et al. Inherited and de novo SHANK2 variants associated with autism spectrum disorder impair neuronal morphogenesis and physiology. Hum Mol Genet 21, 344–357, doi:10.1093/hmg/ddr470 (2012).

29 Zaslavsky, K. et al. SHANK2 mutations associated with autism spectrum disorder cause hyperconnectivity of human neurons. Nat Neurosci 22, 556–564, doi:10.1038/s41593-019-0365-8 (2019).

30 Costas, J. The role of SHANK2 rare variants in schizophrenia susceptibility. Mol Psychiatry 20, 1486, doi:10.1038/mp.2015.119 (2015).

31 Peykov, S. et al. Rare SHANK2 variants in schizophrenia. Mol Psychiatry 20, 1487–1488, doi:10.1038/mp.2015.122 (2015).

32 Won, H. et al. Autistic-like social behaviour in Shank2-mutant mice improved by restoring NMDA receptor function. Nature 486, 261–265, doi:10.1038/nature11208 (2012).

33 Hong, E. J., West, A. E. & Greenberg, M. E. Transcriptional control of cognitive development. Curr Opin Neurobiol 15, 21–28, doi:10.1016/j.conb.2005.01.002 (2005).

34 Nan, X. et al. Interaction between chromatin proteins MECP2 and ATRX is disrupted by mutations that cause inherited mental retardation. Proc Natl Acad Sci U S A 104, 2709–2714, doi:10.1073/pnas.0608056104 (2007).

35 Sullivan, P. F. & Geschwind, D. H. Defining the Genetic, Genomic, Cellular, and Diagnostic Architectures of Psychiatric Disorders. Cell 177, 162–183, doi:10.1016/j.cell.2019.01.015 (2019).

36 Tollervey, J. R. et al. Analysis of alternative splicing associated with aging and neurodegeneration in the human brain. Genome Res 21, 1572–1582, doi:10.1101/gr.122226.111 (2011).

37 Nik, S. & Bowman, T. V. Splicing and neurodegeneration: Insights and mechanisms. Wiley Interdiscip Rev RNA 10, e1532, doi:10.1002/wrna.1532 (2019).

38 Gandal, M. J. et al. Transcriptome-wide isoform-level dysregulation in ASD, schizophrenia, and bipolar disorder. Science 362, doi:10.1126/science.aat8127 (2018).

39 de Leeuw, C. A., Mooij, J. M., Heskes, T. & Posthuma, D. MAGMA: generalized gene-set analysis of GWAS data. PLoS Comput Biol 11, e1004219, doi:10.1371/journal.pcbi.1004219 (2015).

40 Dere, E., Pause, B. M. & Pietrowsky, R. Emotion and episodic memory in neuropsychiatric disorders. Behav Brain Res 215, 162–171, doi:10.1016/j.bbr.2010.03.017 (2010).

41 Guo, J. Y., Ragland, J. D. & Carter, C. S. Memory and cognition in schizophrenia. Mol Psychiatry 24, 633–642, doi:10.1038/s41380-018-0231-1 (2019).

42 Ranganath, C., Minzenberg, M. J. & Ragland, J. D. The cognitive neuroscience of memory function and dysfunction in schizophrenia. Biol Psychiatry 64, 18–25, doi:10.1016/j.biopsych.2008.04.011 (2008).

43 Salmond, C. H. et al. The role of the medial temporal lobe in autistic spectrum disorders. Eur J Neurosci 22, 764–772, doi:10.1111/j.1460-9568.2005.04217.x (2005).

44 Boldog, E. et al. Transcriptomic and morphophysiological evidence for a specialized human cortical GABAergic cell type. Nat Neurosci 21, 1185–1195, doi:10.1038/s41593-018-0205-2 (2018).

45 Mathys, H. et al. Single-cell transcriptomic analysis of Alzheimer’s disease. Nature 570, 332–337, doi:10.1038/s41586-019-1195-2 (2019).

46 Velmeshev, D. et al. Single-cell genomics identifies cell type-specific molecular changes in autism. Science 364, 685–689, doi:10.1126/science.aav8130 (2019).

47 Valnegri, P. et al. The X-linked intellectual disability protein IL1RAPL1 regulates excitatory synapse formation by binding PTPdelta and RhoGAP2. Hum Mol Genet 20, 4797–4809, doi:10.1093/hmg/ddr418 (2011).

48 Um, J. W. & Ko, J. LAR-RPTPs: synaptic adhesion molecules that shape synapse development. Trends Cell Biol 23, 465–475, doi:10.1016/j.tcb.2013.07.004 (2013).

49 Kantojarvi, K. et al. Fine mapping of Xq11.1-q21.33 and mutation screening of RPS6KA6, ZNF711, ACSL4, DLG3, and IL1RAPL2 for autism spectrum disorders (ASD). Autism Res 4, 228–233, doi:10.1002/aur.187 (2011).

50 Lee, K. Y., Jewett, K. A., Chung, H. J. & Tsai, N. P. Loss of fragile X protein FMRP impairs homeostatic synaptic downscaling through tumor suppressor p53 and ubiquitin E3 ligase Nedd4-2. Hum Mol Genet 27, 2805–2816, doi:10.1093/hmg/ddy189 (2018).

51 Zhu, J. et al. Epilepsy-associated gene Nedd4-2 mediates neuronal activity and seizure susceptibility through AMPA receptors. PLoS Genet 13, e1006634, doi:10.1371/journal.pgen.1006634 (2017).

52 Donovan, P. & Poronnik, P. Nedd4 and Nedd4-2: ubiquitin ligases at work in the neuron. Int J Biochem Cell Biol 45, 706–710, doi:10.1016/j.biocel.2012.12.006 (2013).

53 Gao, S. et al. Ubiquitin ligase Nedd4L targets activated Smad2/3 to limit TGF-beta signaling. Mol Cell 36, 457–468, doi:10.1016/j.molcel.2009.09.043 (2009).

54 Kuratomi, G. et al. NEDD4-2 (neural precursor cell expressed, developmentally down-regulated 4-2) negatively regulates TGF-beta (transforming growth factor-beta) signalling by inducing ubiquitin-mediated degradation of Smad2 and TGF-beta type I receptor. Biochem J 386, 461–470, doi:10.1042/BJ20040738 (2005).

55 Chleilat, E. et al. TGF-beta Signaling Regulates Development of Midbrain Dopaminergic and Hindbrain Serotonergic Neuron Subgroups. Neuroscience 381, 124–137, doi:10.1016/j.neuroscience.2018.04.019 (2018).

56 Krieglstein, K., Miyazono, K., ten Dijke, P. & Unsicker, K. TGF-beta in aging and disease. Cell Tissue Res 347, 5–9, doi:10.1007/s00441-011-1278-3 (2012).

57 Krieglstein, K., Zheng, F., Unsicker, K. & Alzheimer, C. More than being protective: functional roles for TGF-beta/activin signaling pathways at central synapses. Trends Neurosci 34, 421–429, doi:10.1016/j.tins.2011.06.002 (2011).

58 Lacmann, A., Hess, D., Gohla, G., Roussa, E. & Krieglstein, K. Activity-dependent release of transforming growth factor-beta in a neuronal network in vitro. Neuroscience 150, 647–657, doi:10.1016/j.neuroscience.2007.09.046 (2007).

59 Fazio, L. et al. Transcriptomic context of DRD1 is associated with prefrontal activity and behavior during working memory. Proc Natl Acad Sci U S A 115, 5582–5587, doi:10.1073/pnas.1717135115 (2018).

60 Zhang, Y. et al. Polymorphisms in human dopamine D2 receptor gene affect gene expression, splicing, and neuronal activity during working memory. Proc Natl Acad Sci U S A 104, 20552–20557, doi:10.1073/pnas.0707106104 (2007).

61 Papassotiropoulos, A. et al. Common Kibra alleles are associated with human memory performance. Science 314, 475–478, doi:10.1126/science.1129837 (2006).

62 Jacobs, J. & Kahana, M. J. Direct brain recordings fuel advances in cognitive electrophysiology. Trends Cogn Sci 14, 162–171, doi:10.1016/j.tics.2010.01.005 (2010).

63 Yamazaki, Y. et al. Short- and long-term functional plasticity of white matter induced by oligodendrocyte depolarization in the hippocampus. Glia 62, 1299–1312, doi:10.1002/glia.22681 (2014).

64 Yamazaki, Y. et al. Region- and Cell Type-Specific Facilitation of Synaptic Function at Destination Synapses Induced by Oligodendrocyte Depolarization. J Neurosci 39, 4036–4050, doi:10.1523/JNEUROSCI.1619-18.2019 (2019).

65 Pepper, R. E., Pitman, K. A., Cullen, C. L. & Young, K. M. How Do Cells of the Oligodendrocyte Lineage Affect Neuronal Circuits to Influence Motor Function, Memory and Mood? Front Cell Neurosci 12, 399, doi:10.3389/fncel.2018.00399 (2018).

66 Yamazaki, Y. et al. Oligodendrocytes: facilitating axonal conduction by more than myelination. Neuroscientist 16, 11–18, doi:10.1177/1073858409334425 (2010).

67 Buzsaki, G. & Moser, E. I. Memory, navigation and theta rhythm in the hippocampal-entorhinal system. Nat Neurosci 16, 130–138, doi:10.1038/nn.3304 (2013).

68 Jacobs, J. Hippocampal theta oscillations are slower in humans than in rodents: implications for models of spatial navigation and memory. Philos Trans R Soc Lond B Biol Sci 369, 20130304, doi:10.1098/rstb.2013.0304 (2014).

69 Nyhus, E. & Curran, T. Functional role of gamma and theta oscillations in episodic memory. Neurosci Biobehav Rev 34, 1023–1035, doi:10.1016/j.neubiorev.2009.12.014 (2010).

70 Vass, L. K. et al. Oscillations Go the Distance: Low-Frequency Human Hippocampal Oscillations Code Spatial Distance in the Absence of Sensory Cues during Teleportation. Neuron 89, 1180–1186, doi:10.1016/j.neuron.2016.01.045 (2016).

71 Yaffe, R. B. et al. Reinstatement of distributed cortical oscillations occurs with precise spatiotemporal dynamics during successful memory retrieval. Proc Natl Acad Sci U S A 111, 18727–18732, doi:10.1073/pnas.1417017112 (2014).

72 Rutishauser, U., Ross, I. B., Mamelak, A. N. & Schuman, E. M. Human memory strength is predicted by theta-frequency phase-locking of single neurons. Nature 464, 903–907, doi:10.1038/nature08860 (2010).

73 Duzel, E., Penny, W. D. & Burgess, N. Brain oscillations and memory. Curr Opin Neurobiol 20, 143–149, doi:10.1016/j.conb.2010.01.004 (2010).

74 Basar, E., Basar-Eroglu, C., Karakas, S. & Schurmann, M. Brain oscillations in perception and memory. Int J Psychophysiol 35, 95–124, doi:10.1016/s0167-8760(99)00047-1 (2000).

75 Kahana, M. J. The cognitive correlates of human brain oscillations. J Neurosci 26, 1669–1672, doi:10.1523/JNEUROSCI.3737-05c.2006 (2006).

76 Vaz, A. P., Inati, S. K., Brunel, N. & Zaghloul, K. A. Coupled ripple oscillations between the medial temporal lobe and neocortex retrieve human memory. Science 363, 975–978, doi:10.1126/science.aau8956 (2019).

77 Lega, B., Burke, J., Jacobs, J. & Kahana, M. J. Slow-Theta-to-Gamma Phase-Amplitude Coupling in Human Hippocampus Supports the Formation of New Episodic Memories. Cereb Cortex 26, 268–278, doi:10.1093/cercor/bhu232 (2016).

78 Ghose, S., Gleason, K. A., Potts, B. W., Lewis-Amezcua, K. & Tamminga, C. A. Differential expression of metabotropic glutamate receptors 2 and 3 in schizophrenia: a mechanism for antipsychotic drug action? Am J Psychiatry 166, 812–820, doi:10.1176/appi.ajp.2009.08091445 (2009).

79 Takahashi, J. S. et al. ChIP-seq and RNA-seq methods to study circadian control of transcription in mammals. Methods in enzymology 551, 285–321, doi:10.1016/bs.mie.2014.10.059 (2015).

80 Habib, N. et al. Massively parallel single-nucleus RNA-seq with DroNc-seq. Nat Methods 14, 955–958, doi:10.1038/nmeth.4407 (2017).

81 Zheng, G. X. et al. Massively parallel digital transcriptional profiling of single cells. Nat Commun 8, 14049, doi:10.1038/ncomms14049 (2017).

82 Dobin, A. et al. STAR: ultrafast universal RNA-seq aligner. Bioinformatics 29, 15–21, doi:10.1093/bioinformatics/bts635 (2013).

83 Wang, L., Wang, S. & Li, W. RSeQC: quality control of RNA-seq experiments. Bioinformatics 28, 2184–2185, doi:10.1093/bioinformatics/bts356 (2012).

84 Anders, S., Pyl, P. T. & Huber, W. HTSeq--a Python framework to work with high-throughput sequencing data. Bioinformatics 31, 166–169, doi:10.1093/bioinformatics/btu638 (2015).

85 Robinson, M. D., McCarthy, D. J. & Smyth, G. K. edgeR: a Bioconductor package for differential expression analysis of digital gene expression data. Bioinformatics 26, 139–140, doi:10.1093/bioinformatics/btp616 (2010).

86 Smith, T., Heger, A. & Sudbery, I. UMI-tools: modeling sequencing errors in Unique Molecular Identifiers to improve quantification accuracy. Genome Res 27, 491–499, doi:10.1101/gr.209601.116 (2017).

87 Liao, Y., Smyth, G. K. & Shi, W. featureCounts: an efficient general purpose program for assigning sequence reads to genomic features. Bioinformatics 30, 923–930, doi:10.1093/bioinformatics/btt656 (2014).

88 Stuart, T. et al. Comprehensive Integration of Single-Cell Data. Cell 177, 1888–1902 e1821, doi:10.1016/j.cell.2019.05.031 (2019).

89 Zhang, Y. et al. Model-based analysis of ChIP-Seq (MACS). Genome Biol 9, R137, doi:10.1186/gb-2008-9-9-r137 (2008).

90 Weirauch, M. T. et al. Determination and inference of eukaryotic transcription factor sequence specificity. Cell 158, 1431–1443, doi:10.1016/j.cell.2014.08.009 (2014).

91 Robinson, J. T., et al. Integrative genomics viewer. Nat Biotechnol 29, 24–26, doi:10.1038/nbt.1754 (2011).

92 Chen, J., Bardes, E. E., Aronow, B. J. & Jegga, A. G. ToppGene Suite for gene list enrichment analysis and candidate gene prioritization. Nucleic Acids Res 37, W305–311 (2009).

93 Grove, J. et al. Identification of common genetic risk variants for autism spectrum disorder. Nat Genet 51, 431–444, doi:10.1038/s41588-019-0344-8 (2019).

94 Jansen, I. E. et al. Genome-wide meta-analysis identifies new loci and functional pathways influencing Alzheimer’s disease risk. Nat Genet 51, 404–413, doi:10.1038/s41588-018-0311-9 (2019).

95 International League Against Epilepsy Consortium on Complex, E. Genome-wide mega-analysis identifies 16 loci and highlights diverse biological mechanisms in the common epilepsies. Nat Commun 9, 5269, doi:10.1038/s41467-018-07524-z (2018).

96 Lee, J. J. et al. Gene discovery and polygenic prediction from a genome-wide association study of educational attainment in 1.1 million individuals. Nat Genet 50, 1112–1121, doi:10.1038/s41588-018-0147-3 (2018).

97 Savage, J. E. et al. Genome-wide association meta-analysis in 269,867 individuals identifies new genetic and functional links to intelligence. Nat Genet 50, 912–919, doi:10.1038/s41588-018-0152-6 (2018).

98 Bipolar, D., Schizophrenia Working Group of the Psychiatric Genomics Consortium. Electronic address, d. r. v. e., Bipolar, D. & Schizophrenia Working Group of the Psychiatric Genomics, C. Genomic Dissection of Bipolar Disorder and Schizophrenia, Including 28 Subphenotypes. Cell 173, 1705–1715 e1716, doi:10.1016/j.cell.2018.05.046 (2018).

99 Davies, G. et al. Study of 300,486 individuals identifies 148 independent genetic loci influencing general cognitive function. Nat Commun 9, 2098, doi:10.1038/s41467-018-04362-x (2018).

100 Wray, N. R. et al. Genome-wide association analyses identify 44 risk variants and refine the genetic architecture of major depression. Nat Genet 50, 668–681, doi:10.1038/s41588-018-0090-3 (2018).

101 Pardinas, A. F. et al. Common schizophrenia alleles are enriched in mutation-intolerant genes and in regions under strong background selection. Nat Genet 50, 381–389, doi:10.1038/s41588-018-0059-2 (2018).

102 Martin, J. et al. A Genetic Investigation of Sex Bias in the Prevalence of Attention-Deficit/Hyperactivity Disorder. Biol Psychiatry 83, 1044–1053, doi:10.1016/j.biopsych.2017.11.026 (2018).

103 Sohail, M. et al. Polygenic adaptation on height is overestimated due to uncorrected stratification in genome-wide association studies. Elife 8, doi:10.7554/eLife.39702 (2019).

104 Hoffmann, T. J. et al. A Large Multiethnic Genome-Wide Association Study of Adult Body Mass Index Identifies Novel Loci. Genetics 210, 499–515, doi:10.1534/genetics.118.301479 (2018).

105 Morris, A. P. et al. Large-scale association analysis provides insights into the genetic architecture and pathophysiology of type 2 diabetes. Nat Genet 44, 981–990, doi:10.1038/ng.2383 (2012).

106 Estrada, K. et al. Genome-wide meta-analysis identifies 56 bone mineral density loci and reveals 14 loci associated with risk of fracture. Nat Genet 44, 491–501, doi:10.1038/ng.2249 (2012).

107 Schunkert, H. et al. Large-scale association analysis identifies 13 new susceptibility loci for coronary artery disease. Nat Genet 43, 333–338, doi:10.1038/ng.784 (2011).

